# The dominance hierarchy of the female Yunnan snub-nosed monkeys (*Rhinopithecus bieti*)

**DOI:** 10.1101/2020.03.19.998781

**Authors:** Kai Huang, Wancai Xia, Yi Fu, Yaqiong Wan, Hao Feng, Ali Krzton, Jiaqi Li, Dayong Li

## Abstract

Dominance hierarchies are common in social mammals, especially primates. The formation of social hierarchies is conducive to solving the problem of the allocation of scarce resources among individuals. From August 2015 to July 2016, we observed a wild, provisioned Yunnan snub-nosed monkey (*Rhinopithecus bieti*) group at Xiangguqing in Baimaxueshan National Nature Reserve, Yunnan Province, China. Aggressive and submissive behaviors were used to investigate dominance hierarchies between female individuals in the same one-male unit (OMU), and the grooming reciprocity index was used to detect reciprocal relationships between these females within the OMU. The results showed that loose social hierarchies exist among the females in each OMU, and more dominant individuals have higher grooming incomes. These results are consistent with the aggressive-submissive hypothesis and the resource control hypothesis.

## Introduction

Dominance hierarchies are one outcome of the long-term evolution of social animals [1]. Hierarchy is common in social mammals, especially the primates [2]. Recently, primate dominance hierarchies have become a hot topic in behavioral ecology, including studies of *Rhinopithecus roxellana, Macaca thibetana, M. fuscata, M. assamensis, Pan paniscus, Papio hamadryas* and *Saimiri sciureus* [3]. Individuals may form dominance hierarchies in order to stabilize the community structure, optimize the use of natural resources, avoid casualties from fights between individuals, and protect weaker group members so as to better adapt to the ecological conditions they experience [4]. A dominance hierarchy refers to a ranked order of individuals within a social group as reflected by certain behavioral patterns. Dominance hierarchies can allow certain individuals to control desirable resources and maintain unit stability [5]. Within the primate order, dominance hierarchies have been significant adaptations to species’ gregarious lifestyles [6]. On this basis, some researchers proposed the aggressive-submissive hypothesis and the priority occupy resources hypothesis to explain the biological function and ecological significance of the reproductive unit dominance hierarchy of primate [7].

The aggressive-submissive hypothesis states that the dominance hierarchy is formed according to the relative superiority of senders and receivers of inter-individual aggressive and submissive behaviors [8,9]. The dominance hierarchy within a group develops after repeated competition between individuals and can be described according to the asymmetric aggression and submission patterns that occur [7]. Wang et al (2007) studied the relationship between dominance hierarchies and food competition in Sichuan snub-nosed monkeys (*Rhinopithecus roxellana*), and those results were consistent with this hypothesis [10]. When environmental resources are insufficient to satisfy all individuals in the group, competition will occur [11]. Over the long term, repeated bouts of aggression and submission promote stable dominance hierarchies, making it easier for higher-ranking individuals to obtain more and better resources [12]. The aggressive-submissive hypothesis has empirical support in some primate species [13-15].

The resource control hypothesis suggests that higher-ranking individuals can secure better access to limited, desirable resources [3,9]. Research on dominance hierarchies in *R. roxellana* and *Cebus capucinus* have shown that higher-ranking group members claim safer spaces and better food before others [3,16]. Dominance hierarchies can substantially affect the quality of life and reproductive success of female primates [17]. The female dominance hierarchy within a group, as reflected in the frequency and intensity of social interactions among individuals, can determine access to high-quality resources and mates [18]. In polygynous primates with female dominance hierarchies, high-ranking females are more likely to be favored by unit males, and they may also restrict mating opportunities for lower-ranking females [19,20,21,22].

So far, there are no published studies of female dominance hierarchies in wild populations of *R. biet*i [9,11], but studying such hierarchies within the one-male units (OMUs) that comprise *R. bieti*’s multilevel society will improve our understanding of community structure and stability in this species. This is conducive to the protection and scientific management of this rare and endangered species [23,24]. In this study, reliable individual identification of monkeys allowed the female dominance hierarchies within the study group to be described on the basis of bouts of aggressive and submissive behavior. Exploring the relationship between female dominance hierarchies and grooming income permitted the testing of the aggressive-submissive hypothesis and the resource control hypothesis.

## Materials and methods

### Behavioral definitions and data collection

The frequencies of aggressive and submissive behavior on the part of individual females were used to map dominance hierarchies within each one-male unit. According to Li et al. (2006), the following aggressive behaviors are characteristic of *R. bieti*: bite, snatch, catch, threat, and displacement; submissive behaviors were divided into avoidance, crouch, and escape [8]. From August 2015 to July 2016, the effective observation time was 231 days. Aggressive and submissive behavior was observed and recorded for named individuals using focal animal sampling and all-occurrence sampling [25]. For each occurrence we recorded the initiator, the recipient, the type of conflict behavior, and the cause and effect of the conflict [8].

### Data analyses

We calculated values for DS and NDS to determine the dominance hierarchy of females within an OMU [26]. 

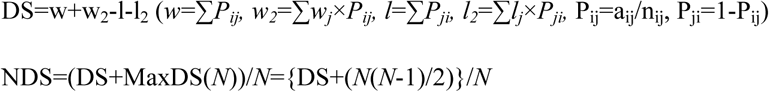

where *w* represents the sum of *i*’s *P*_*ij*_ values, *w*_*2*_ represents the summed *w* values (weighted by the appropriate *P*_*ij*_ values). *l* represents the sum of *i*’s *P*_*ji*_ values and *l*_*2*_ represents the summed *l* values (weighted by the appropriate *P*_*ji*_ values) of those individuals with which *i* interacted. *P*_*ij*_ represents the proportion of wins by individual *i* in her interactions with another individual *j. a*_*ij*_ represents the number of times *i* individuals defeat *j* individuals, *n*_*ij*_ represents the total number of interactions between *i* and *j*. If there is no aggression or submission between individual *i* and individual *j*, then *P*_*ij*_=*P*_*ji*_=0. MaxDS indicates the maximum possible DS of the individual *i*, while *N* represents the number of individuals.

The grooming reciprocity index was used to measure the degree of reciprocity in each female dyad in an OMU [27]. 

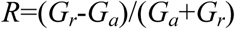

*R* represents the grooming reciprocity index, *G*_*a*_ represents the grooming time initiated by individual A, and *G*_*r*_ represents the grooming time received by individual A. Smaller values of *R* signify less benefits from grooming.

The dominance hierarchy is the absolute value of the slope of the curve plotted with values of DS on the x-axis and NDS on the y-axis and always falls between 0 and 1. A dominance hierarchy value close to 1 indicates a group that is rigidly stratified, whereas a group with a value closer to 0 is more egalitarian. Linear regression and the Spearman correlation were calculated to test the relationship between the female dominance hierarchies and grooming benefits. All collected data was processed and analyzed using Microsoft Excel 2013 and SPSS 22.0, with significance levels set to 0.05.

## Results

### Female dominance hierarchy within each OMU

Aggressive and submissive behavioral data was recorded for five OMUs; three other OMUs (Duanshou, Danba, and Lianheguo) were excluded on the basis of having only one female during the study period.

A total of 2,947 aggressive or submissive behaviors were observed, of which 2,545 occurred between adult females within the OMU. The remaining 402 behaviors, which occurred between members of different OMUs, were not analyzed. The number of aggressive or submissive behaviors recorded for each focal OMU were as follows: HD - 873, DGZ - 588, HL - 617, LB - 402, and XS - 449 (Table 1).

**Table 1.**
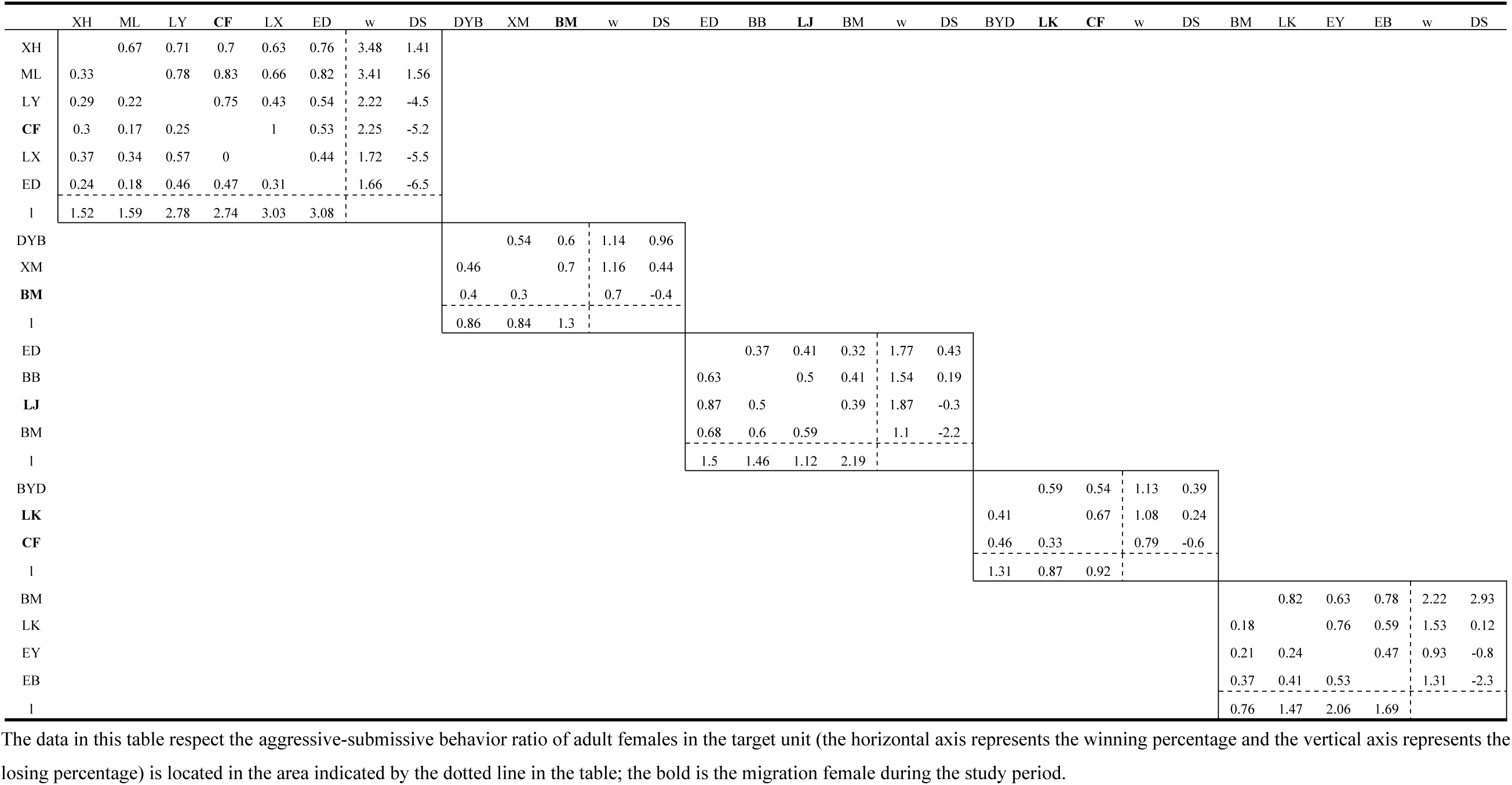
The female dominance hierarchies among the five focal OMUs.

The results showed that each OMU had a stable female dominance hierarchy: XH> ML> LY> CF> LX> ED for the HD OMU, DYB> XM> BM for the DGZ OMU, ED> BB> LJ> BM for the HL OMU, BYD> LK> CF for the LB OMU, and BM> LK> EY> EB for the XS OMU. On the other hand, these dominance hierarchies were not strict, with values closer to 0 than to 1 (*K*_*s* (HD)_ = 0.29; *K*_*s*(DGZ)_=0.23; *K*_*s*(HL)_=0.21; *K*_*s*(LB)_=0.17; *K*_*s*(XS)_=0.41) (Fig. 1).

**Fig. 1.**
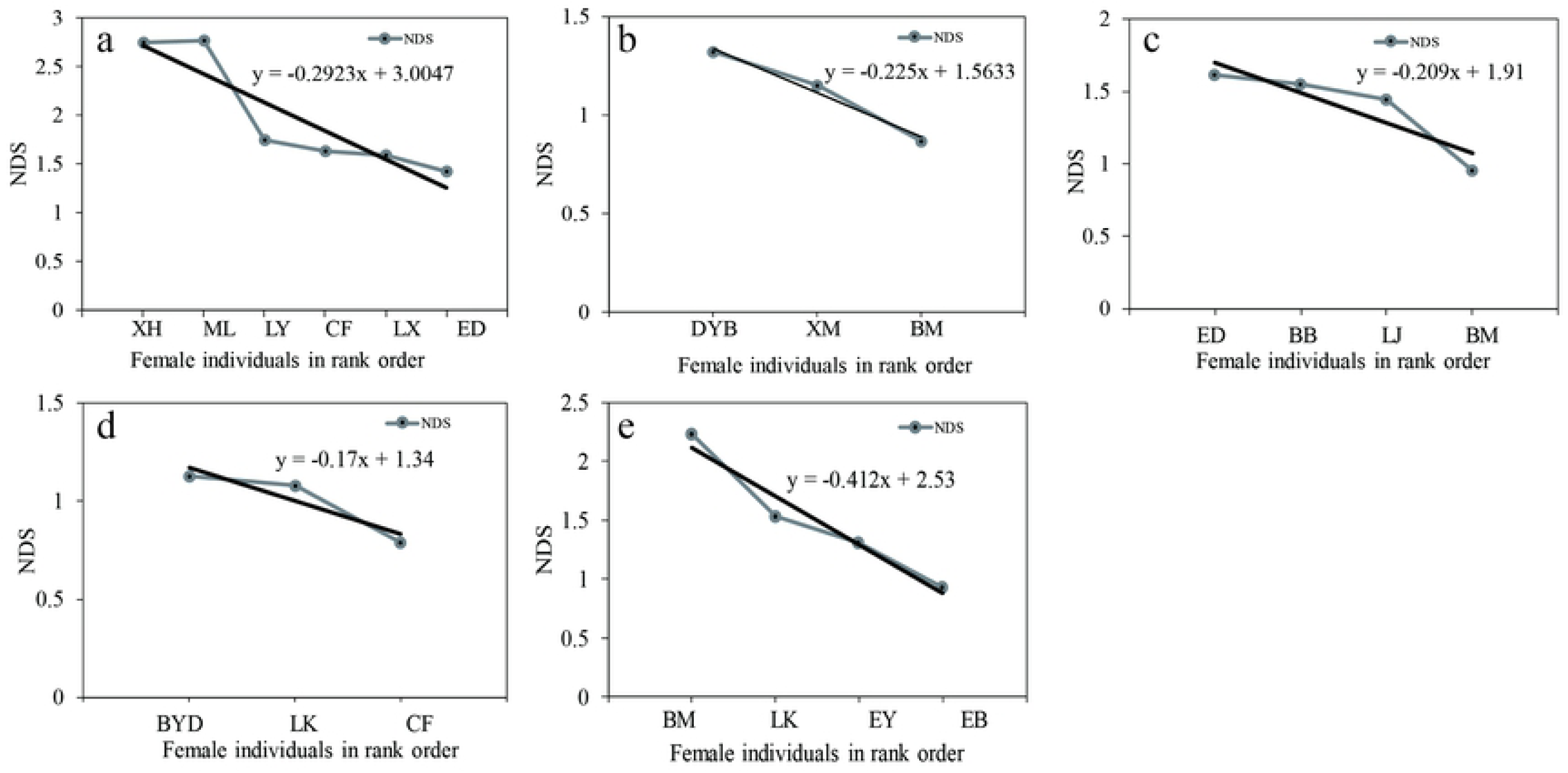
Female hierarchical steepness during the study period (a: HD OMU; b: DGZ OMU; c: HL OMU; d: LB OMU; e: XS OMU)

### Dominance hierarchy and grooming reciprocity

A total of 3,788.92 minutes of grooming was recorded between females during the study period (by OMU: HD -1272.22min, DGZ -665.3min, HL -881.18min, LB -574.57min, XS -325.65min). Since some females migrated into or out of the focal OMUs during the study period, grooming time was normalized using the ratio (time initiating or receiving grooming/time of individual observation). The analysis showed that higher-ranking females had higher grooming incomes, with each OMU having a significantly positive Spearman correlation between female rank and grooming income (Spearman correlation test: *R*_*HD*_ =0.886, *P*<0.05; *R*_*DGZ*_ =0.999, *P*<0.05; *R*_*HL*_ =0.981, *P*<0.05; *R*_*LB*_ =0.998, *P*<0.05; *R*_*XS*_ =0.98, *P*<0.05) (Fig. 2).

**Fig. 2.**
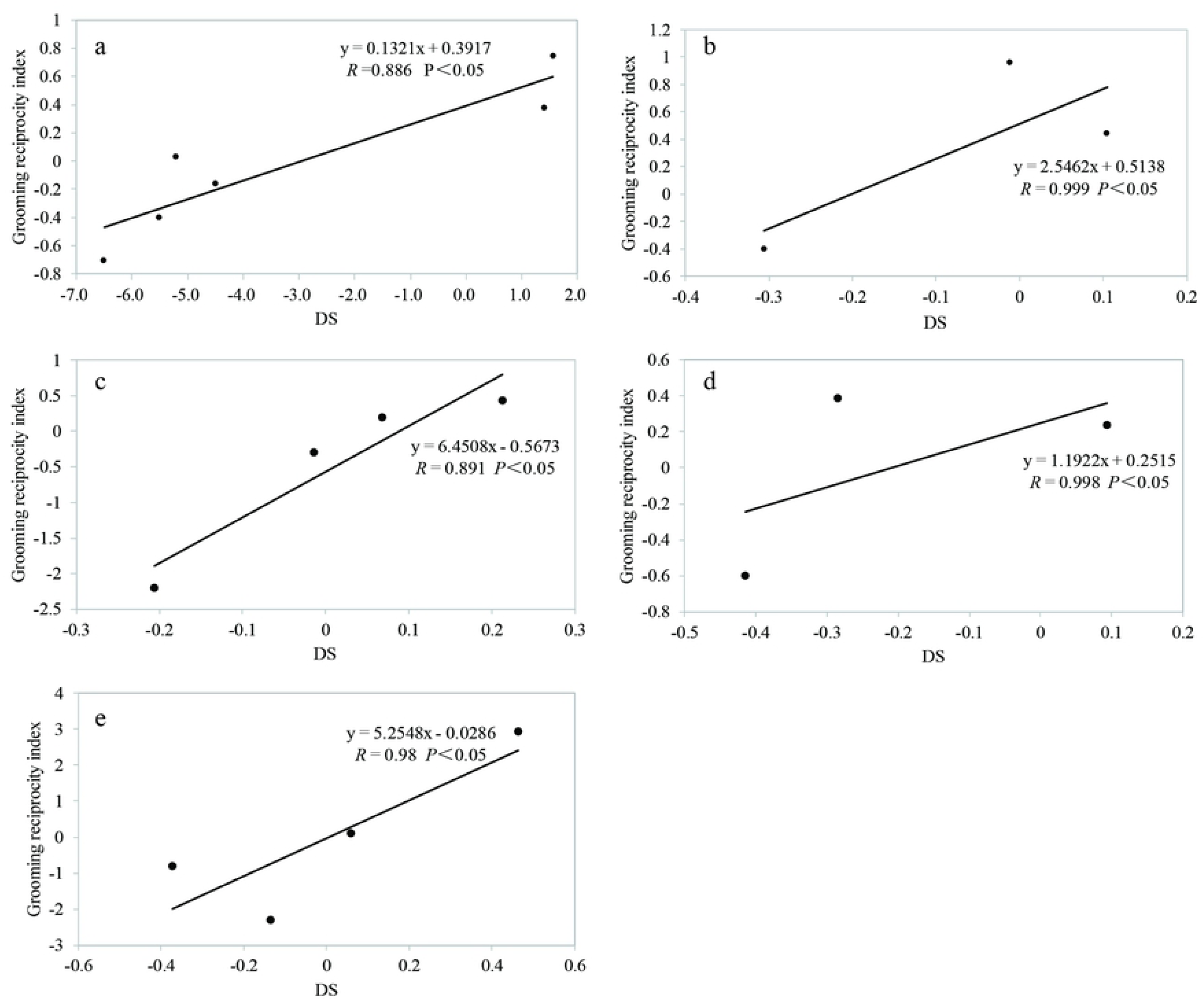
Correlation between female dominance hierarchy and grooming reciprocity (a: HD OMU; b: DGZ OMU; c: HL OMU; d: LB OMU; e: XS OMU)

## Discussion

### The Aggressive-submissive Hypothesis

Adult female *R. bieti* rarely fight each other. Yu et al. (2009) have shown that in the congeneric Sichuan snub-nosed monkey (*R. roxellana*), fighting behavior only occured between males at Shennongjia [28]. Studies on captive Yunnan snub-nosed monkeys have suggested that *R. bieti* is a relatively tolerant species, with hierarchies that are correspondingly loose [11]. Moreover, high-ranking individuals do not monopolize quality resources, giving individuals of lower rank the opportunity to obtain a share [29]. Females deal with competition through ritualized behavior, which can determine the dominance hierarchy without the need for direct conflict and allow individuals to avoid injury [8]. At the same time, reproductive costs are much higher for females than males, and their reproductive success is tightly constrained by nutritional stress. Therefore, females are competing indirectly for resources as opposed to males competing directly for mating opportunities.

Female Yunnan snub-nosed monkeys appear to substitute low-risk exchanges of dominance and submission for overt aggressive behavior, which is consistent with the aggressive-submissive hypothesis. It has been argued that if attacks between individuals are rare, a dominance hierarchy does not exist or is implicit [30,31]. However, this study confirmed that relationships between adult females in this species are still hierarchical.

Similar results have been found in studies of *Pan troglodytes* [32,33,34] and *R. roxellana* [35]. The dominance hierarchy manifests after repeated asymmetric agonistic interactions between individuals, which may be limited to certain contexts that depend on environmental conditions [7]. For instance, the distribution and relative scarcity in space and time can affect individuals’ mutual tolerance. *R.bieti* mainly feed on low-energy lichens, which are evenly distributed and consistently available, for most of the year. Scarce, high-quality foods such as the fruit of *Cornus capitata* appear only at certain times of year, which could increase aggression and reduce tolerance during that season as individuals scramble to consume those foods.

### The Resource Control Hypothesis

The balance of giving vs. receiving in bouts of allogrooming can reflect the dominance hierarchy. It has been argued that in some species, such as the capuchin monkey (*Cebus apella*), allogrooming is an expression of mutual attraction, and its frequency is unaffected by rank or kinship status [36]. However, our findings for grooming income in *R. bieti* were similar to patterns observed in green monkeys (*Chlorocebus sabaeus*), bonobos (*Pan paniscus*), and stump-tailed macaques (*Macaca arctoides*) [37,38,39]. Allogrooming behavior has a social function, reducing tension and regulating hierarchical relationships between group members such that higher-ranking individuals receive more grooming [40]. Allogrooming may serve as a ritualized way to reduce conflict, as low-ranking individuals might groom higher-ranking individuals in exchange for tolerance and reduced aggression [40,41]. In wild *R. roxellana*, for instance, individuals that are attacked or threatened will groom the aggressor after the conflict has subsided [42]. Adult female collared mangabeys (*Cercocebus torquatus lunulatus*) that are attacked by another group member will attempt reconciliation via grooming to avoid being attacked again [43], and in bonnet macaques (*M. radiata*), low-ranking individuals are not attacked by other group members while grooming high-ranking individuals [44]. In short, a grooming economy can emerge when low-ranking individuals trade grooming for reduced conflict and protection, allowing high-ranking individuals to gain more grooming income [45,46].

## Conclusions

In conclusion, we presented the first detailed analysis of dominance hierarchies in wild adult females within the same OMU in *R. bieti*. We argue that these females do have hierarchical relationships, but that they substitute regular displays of aggression and submission for costly and potentially dangerous direct conflicts, consistent with the aggressive-submissive hypothesis. Similarly to other primates, allogrooming behavior in *R. bieti* functions to reduce tension and reinforce the dominance hierarchy, allowing high-ranking individuals to benefit more from grooming. Grooming can also be used as a means of quelling conflicts or mitigating attacks, consistent with the resource control hypothesis.

## Acknowledgments

We are grateful to our field assistants Jianhua Yu, Jinming Yu and Lizhong Yu. We thank Baimaxueshan National Nature Reserve for our work permit. This research was funded by the National Key Program of Research and Development, Ministry of Science and Technology (No. 2016YFC0503200; 2017YFC0505205), the Second Tibetan Plateau Scientific Expedition and Research Program (No. 2019QZKK0501), National Natural Science Foundation of China (No.31470461), Applied Basic Research Program of Sichuan Province (No. 2017JY0325), the Biodiversity Investigation, Observation and Assessment Program of Ministry of Ecology and Environment of China.

